# On the Complexity of Sequence to Graph Alignment

**DOI:** 10.1101/522912

**Authors:** Chirag Jain, Haowen Zhang, Yu Gao, Srinivas Aluru

**Author notes:** Should be regarded as joint first-authors (Chirag Jain, Haowen Zhang).

## Abstract

Availability of extensive genetics data across multiple individuals and populations is driving the growing importance of graph based reference representations. Aligning sequences to graphs is a fundamental operation on several types of sequence graphs (variation graphs, assembly graphs, pan-genomes, etc.) and their biological applications. Though research on sequence to graph alignments is nascent, it can draw from related work on pattern matching in hypertext. In this paper, we study sequence to graph alignment problems under Hamming and edit distance models, and linear and affine gap penalty functions, for multiple variants of the problem that allow changes in query alone, graph alone, or in both. We prove that when changes are permitted in graphs either standalone or in conjunction with changes in the query, the sequence to graph alignment problem is 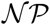-complete under both Hamming and edit distance models for alphabets of size ≥ 2. For the case where only changes to the sequence are permitted, we present an *O*(|*V*| + *m*|*E*|) time algorithm, where *m* denotes the query size, and *V* and *E* denote the vertex and edge sets of the graph, respectively. Our result is generalizable to both linear and affine gap penalty functions, and improves upon the run-time complexity of existing algorithms.

## 1 Introduction

Aligning sequences to graphs is becoming increasingly important in the context of several applications in computational biology, including variant calling [22, 6, 7, 9], genome assembly [2, 33, 8], long read error-correction [28, 32, 34], RNA-seq data analysis [4, 13], and more recently, antimicrobial resistance profiling [27]. Much of this has been driven by the growing ease and ubiquity of sequencing at personal, population, and environmental-scale, leading to significant growth in availability of datasets. Graph based representations provide a natural mechanism for compact representation of related sequences and variations among them. Some of the most useful graph based data structures are de-Bruijn graphs [25], variation graphs [23], string graphs [19], and partial order graphs [14].

Decades of progress made towards designing provably good algorithms for the classic sequence to sequence alignment problems serves as the foundation for mapping tools currently used in genomics, and similar efforts are necessary for sequence to graph alignment. To address the growing list of biological applications that require aligning sequences to a graph, several heuristics [12, 16, 15, 11, 9] and a few provably good algorithms [29, 26, 31] have been developed in recent years. In addition, sequence to graph alignment has been studied much earlier in the string literature through its counterpart, approximate pattern matching to hypertext [17]. Since then, important complexity results and algorithms have been obtained for different variants of this problem [1, 20, 30].

Many versions of the classic sequence to sequence alignment problem were considered in the literature, e.g., different alignment modes – local/global, scoring functions – linear/affine/arbitrary gap penalty, and so on [21]. The list further proliferates when considering a graph-based reference. This is because the nature of the problem changes depending on whether the input graphs are cyclic or acyclic [20], and whether edits are allowed in the graph, or query, or both [1].

In this paper, we present new complexity results and improved algorithms for multiple variants of the sequence to graph alignment problem. Consider a query sequence of length *m* and a directed graph *G*(*V*, *E*) with string-labeled vertices, over the alphabet *Σ*. We make the following contributions:

- The problem variants that allow changes to the graph labels are shown to be 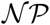-complete [1], via proofs that assume |*Σ*| ≥ |*V*|. To date, tractability of these problems remains unknown for the case of constant sized alphabets, which is an important consideration when aligning DNA, RNA, or protein sequences to corresponding graphs. We close this knowledge gap by proving that four variants of the problem, characterized by changes to graph alone or both graph and query, under the Hamming or edit distance models, remain 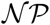-complete for |*Σ*| ≥ 2.
- Allowing changes to the query sequence alone makes the problem polynomially solvable. For graphs with character-labeled vertices, we propose an algorithm that achieves *O*(|*V*| + *m*|*E*|) time bound for both linear and affine gap penalty cases, superior to the best existing algorithms (Table 1). An important attribute of the proposed algorithm is that it achieves the same time and space complexity as required for the easier problem of sequence alignment to acyclic graphs [17, 20], under both scoring models.

## 2 Preliminaries

Let *Σ* denote an alphabet, and *x* and *y* be two strings over *Σ*. We use *x*[*i*] to denote the *i*^*th*^ character of *x*, and |*x*| to denote its length. Let *x*[*i*, *j*] (1 ≤ *i* ≤ *j* ≤ |*x*|) denote *x*[*i*]*x*[*i* + 1] … *x*[*j*], the substring of *x* beginning at the *i*^*th*^ position and ending at the *j*^*th*^ position. Concatenation of *x* and *y* is denoted as *xy*. Let *x*^*k*^ denote string *x* concatenated with itself *k* times.

### Definition 1.

*Sequence Graph: A sequence graph G*(*V*, *E*, *σ*) *is a directed graph with vertices V and edges E. Function σ* : *V → Σ* labels each vertex v* ∈ *V with string σ*(*v*) *over the alphabet Σ.*

Naturally, path *p* = *v*_*i*_, *v*_*i*+1_, …, *v*_*j*_ in *G*(*V*, *E*, *σ*) spells the sequence *σ*(*v*_*i*_) *σ*(*v*_*i*+1_) … *σ*(*v*_*j*_). Given a query sequence *q*, we seek its best matching path sequence in the graph. Alignment problems are formulated such that distance between the computed path and the query sequence is minimized, subject to a specified distance metric such as Hamming or edit distance. Typically, an alignment is scored using either a linear or an affine gap penalty function. The cost of a gap is proportional to its length, when using a linear gap penalty function. An affine gap penalty function imposes an additional constant cost to initiate a gap.

**Table 1.** Comparison of run-time complexity achieved by different algorithms for the sequence to graph alignment problem when changes are allowed in the query sequence alone.

|  | Linear gap penalty |  | Affine gap penalty |
| --- | --- | --- | --- |
|  | Edit distance | Arbitrary costs |  |
| Amir *et al.* [1] | $O(m(\vert V\vert \log \vert V\vert + \vert E\vert ))$ | $O(m(\vert V\vert \log \vert V\vert + \vert E\vert ))$ | - |
| Navarro [20] | $O(m(\vert V\vert + \vert E\vert ))$ | - | - |
| HybridSpades [2] | $O(m(\vert V\vert \log(m\vert V\vert ) + \vert E\vert ))$ | $O(m(\vert V\vert \log(m\vert V\vert ) + \vert E\vert ))$ | - |
| V-ALIGN [31] | $O(m\vert V\vert \vert E\vert )$ | $O(m\vert V\vert \vert E\vert )$ | $O(m\vert V\vert \vert E\vert )$ |
| Rautiainen and Marschall [26] | $O(\vert V\vert + m\vert E\vert )$ | $O(m(\vert V\vert \log \vert V\vert + \vert E\vert ))$ | $O(m(\vert V\vert \log \vert V\vert + \vert E\vert ))$ |
| **This work** | $O(\vert V\vert + m\vert E\vert )$ | $O(\vert V\vert + m\vert E\vert )$ | $O(\vert V\vert + m\vert E\vert )$ |

## 3 Complexity Analysis

### 3.1 Asymmetry of Edit Locations

An alignment between two sequences also specifies possible changes to the sequences (e.g. substitutions, insertions, deletions) to make them identical, with alignment distance specifying the cumulative penalty for the changes. The changes can be individually applied either to the first or the second sequence, or any combination thereof. Such a symmetry is no longer valid when aligning sequences to graphs [1]. This is because alignments can occur along cyclic paths in the graph. If the label of a vertex in the graph is changed, then an alignment path visiting that vertex *k* times reflects the same change at *k* different positions in the alignment. On the other hand, a change in one position of the sequence only reflects that change in the corresponding position in the alignment. As such, optimal alignment scores vary depending on whether changes are permitted in just the sequence, just the graph, or both (see Figure 1 for an illustration). This characteristic leads to *three different problems*, with each potentially resulting in a different optimal distance.

**Fig. 1.**
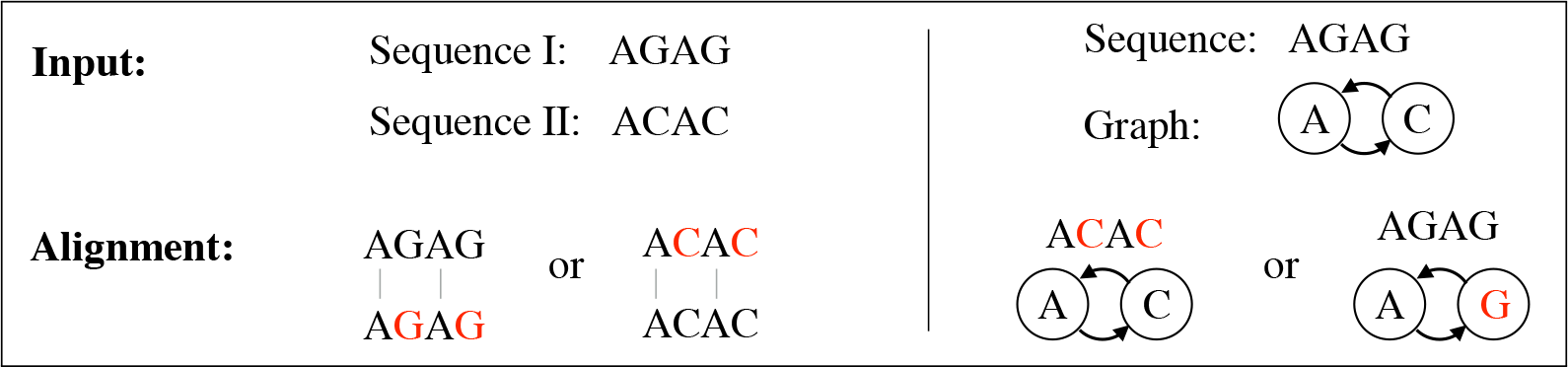
Asymmetry w.r.t. the location of changes in sequence to graph alignment illustrated using Hamming distance. Two substitutions are required in the sequence, whereas just one is sufficient if made in the graph.

Consider the sequence to graph alignment problem under the Hamming or edit distance metrics. For each distance metric, there are three versions of the problem depending on whether changes are allowed in query alone, graph alone, or both in the query and graph. Consider the decision versions of these problems, which ask whether there exists an alignment with ≤ *d* modifications (substitutions or edits), as per the distance metric. Restricting substitutions or edits to the query sequence alone admits polynomial time solutions [1, 20, 26]. In the pioneering work of Amir *et al.* [1] in the domain of string to hypertext matching, it has been proved that the other problem variants which permit changes to graph are 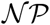-complete. The proofs provided in their work assume an alphabet size ≥ |*V*|. To date, tractability of these problems remains unknown for the case of constant sized alphabets (e.g., for DNA, RNA, or protein sequences). In what follows, we close this knowledge gap by showing that the problems remain 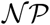-complete for any alphabet of size at least 2.

### 3.2 Alignment using Hamming Distance

#### Theorem 1

*The problem “Can we substitute a total of ≤ d characters in graph G and query q such that q will have a matching path in G?” is 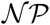-complete for |Σ| ≥ 2*.

*Proof.* The problem is in 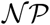. Given a solution, the set of substitutions can be used to obtain the corrected graph and query. Next, we can leverage any polynomial time algorithm [1, 20, 24] to verify if the corrected query matches a path in the corrected graph.

To show that the problem is 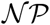-hard, we perform a reduction using the directed Hamiltonian cycle problem. Suppose *G*′(*V*, *E*) is a directed graph in which we seek a Hamiltonian cycle. Let *n* = |*V*|. We transform it into a sequence graph *G*(*V*, *E*, *σ*) over the alphabet *Σ* = {*α*, *β*} by simply labeling each vertex *v* ∈ *V* with *α*^*n*^ (Figure 2). Note that the graph structure remains unchanged. Next, we construct query sequence *q*. Let token *t*_*i*_ be the sequence of *n* characters *α*^*n*−*i*−1^ *βα*^*i*^. We choose query *q* to be the *n*^2^(2*n* + 2) long sequence: (*t*_0_*t*_1_ … *t*_*n*−1_)^2*n*+2^. We claim that a Hamiltonian cycle exists in *G*′(*V*, *E*) if and only if *q* can be matched after substituting a total of ≤ *n* characters in *G*(*V*, *E*, *σ*) and *q*.

**Fig. 2.**
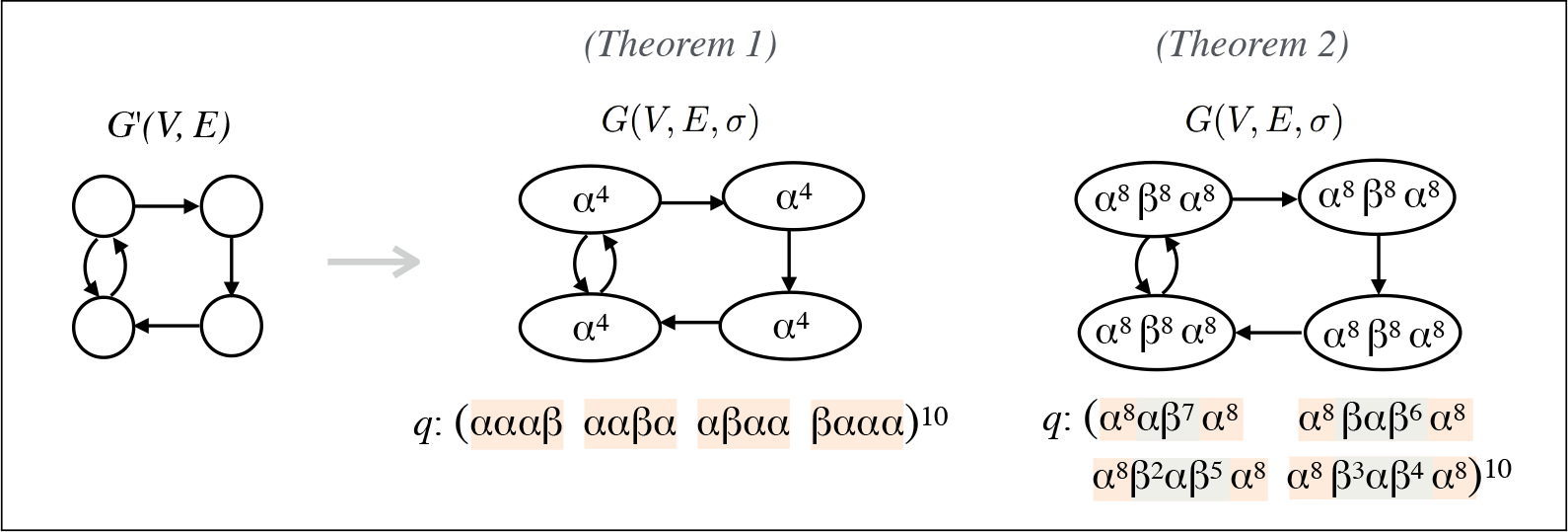
The constructs used for reductions in proofs of Theorems 1 and 2.

Suppose there is a Hamiltonian cycle in *G*′(*V*, *E*). We can follow the corresponding loop in *G*(*V*, *E*, *σ*) from the first character of any vertex label. To match each token in the query *q*, we require one *α* → *β* substitution per vertex. Thus, the query *q* matches *G*(*V*, *E*, *σ*) after making exactly *n* substitutions in the graph.

Conversely, suppose the query *q* matches the graph *G*(*V*, *E*, *σ*) after making ≤ *n* substitutions in the query and the graph. Consider the following substring *q*_*sub*_ of *q*: *t*_0_*t*_1_ … *t*_*n*−1_*t*_0_*t*_1_. Note that there are *n* + 1 non-overlapping instances of *q*_*sub*_ in *q*. Even if all the *n* substitutions occur in the query, at least one instance of *q*_*sub*_ must remain unchanged. As a result, *q*_*sub*_ must match to a path in the corrected *G*(*V*, *E*, *σ*).

#### Case 1

*q_sub_ starts matching from the first character of a vertex label.* Note that the first *n* tokens *q*_*sub*_[1, *n*] = *t*_0_, *q*_*sub*_[*n* + 1, 2*n*] = *t*_1_, …, *q*_*sub*_[*n*^2^ − *n* + 1, *n*^2^] = *t*_*n*−1_ are all unique followed by *q*_*sub*_[*n*^2^ + 1, *n*^2^ + *n*] = *t*_0_. Therefore, this requires a Hamiltonian cycle in *G*(*V*, *E*, *σ*). Accordingly, there is a Hamiltonian cycle in *G*′(*V*, *E*).

#### Case 2

*q_sub_ starts somewhere other than the starting position within a vertex label.* Let *q*_*sub*_[*k*] (1 < *k* ≤ *n*) be the first character that matches at the beginning of the next vertex on the path matching *q*. Similar to the previous case, the following *n* sequences *q*_*sub*_[*k*, *n* + *k* − 1], *q*_*sub*_[*n* + *k*, 2*n* + *k* − 1], …, *q*_*sub*_[*n*^2^ − *n* + *k*, *n*^2^ + *k* − 1] are unique due to the spacing between *β* characters in *q*_*sub*_. Therefore, the matching path must yield a Hamiltonian cycle.

#### Corollary 1

*The problem “Can we substitute ≤ d characters in graph G such that q will have a matching path in G?” is 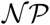-complete for |Σ| ≥* 2.

*Proof.* The setup used in the proof of Theorem 1 can be trivially extended to prove the above claim. Alternatively, we can simplify the proof by using the query sequence *q* = (*t*_0_*t*_1_ … *t*_*n*−1_)^2^ since only one instance of the substring *q*_*sub*_ in *q* is needed for the subsequent arguments. This is because substitutions in the query sequence are not permitted.

Using the above two results, we conclude that Hamming-distance based decision formulations of sequence to graph alignment problems are 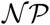-complete when substitutions are allowed in graph labels, for |*Σ*| ≥ 2. In fact, it can be easily shown that |*Σ*| ≥ 2 reflects a tight bound. Using |*Σ*| = 1, all the problem instances can be decided in polynomial time using straightforward application of standard graph algorithms.

### 3.3 Alignment using Edit Distance

We next show that edit distance based decision problems that permit changes in graph labels are 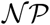-complete if |*Σ*| ≥ 2. Similar to our previous claims, allowing edits in the graph makes the sequence to graph alignment problem intractable. Proofs used for Hamming distance do not apply here as edits also permit insertions and deletions. Length of vertex labels can grow or shrink using insertion and deletion edits respectively.

#### Theorem 2

*The problem “Can we perform a total of ≤ d edits in graph G and query q so that q will match in G?” is 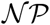-complete for |Σ| ≥* 2.

*Proof.* Clearly the problem is in 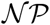. We again use the directed Hamiltonian cycle problem for reduction. Given an instance *G*′(*V*, *E*) of the directed Hamiltonian cycle problem, we design an instance *G*(*V*, *E*, *σ*) using *Σ* = {*α*, *β*}. Let *n* = |*V*|. Label each vertex *v* in *V* using a sequence of 6*n* characters *α*^2*n*^ *β*^2*n*^*α*^2*n*^ (Figure 2). Let token *t*_*i*_ be a sequence of length 6*n*: *α*^2*n*^ *β*^*i*^*αβ*^2*n*−1−*i*^ *α*^2*n*^. Using such tokens, we build a query sequence *q* of length 6*n*^2^(2*n* + 2) as (*t*_0_*t*_1_ … *t*_*n*−1_)^2*n*+2^. We claim that a Hamiltonian cycle exists in *G*′(*V*, *E*) if and only if we can match the sequence *q* to the graph *G*(*V*, *E*, *σ*) using ≤ *n* total edits.

If there is a Hamiltonian cycle in *G*′(*V, E*), we can follow the same loop in *G*(*V*, *E*, *σ*) to align *q*. The alignment requires one substitution per vertex. To prove the converse, suppose query *q* matches graph *G*(*V*, *E*, *σ*) after making a total of ≤ *n* edits in *q* and *G*(*V*, *E*, *σ*). Consider the substring *q*_*sub*_ of *q*: *t*_0_*t*_1_ … *t*_*n*−1_*t*_0_. Note that there are *n* + 1 non-overlapping instances of *q*_*sub*_ in *q*, at least one of which must remain unchanged. Accordingly, the substring *q*_*sub*_ must match corrected *G*(*V*, *E*, *σ*).

For the token *t*_*i*_, let *k*_*i*_ = *β*^*i*^*αβ*^2*n*−1−*i*^ be its *kernel* sequence of length 2*n*. It follows that *t*_*i*_ = *α*^2*n*^*k*_*i*_*α*^2*n*^. We show that a kernel must be matched entirely within a vertex in *G*(*V*, *E*, *σ*) using the following two arguments. First, since any vertex label cannot shrink from length 6*n* to < 5*n*, a kernel cannot be matched to an entire vertex after the edits. It implies that a kernel must match to ≤ 2 vertices. Second, if a kernel aligns across two vertices, (2*n* − 1) *β*’s must be required in place of *α*’s at the two vertex ends, thus requiring > *n* edits. Therefore, a kernel can only be matched within a single vertex label. Finally, it is easy to observe that any vertex label after ≤ *n* edits cannot be matched to more than one kernel. When combining these arguments with the fact that all *n* consecutive kernels in *q*_*sub*_ are unique, we establish that the alignment path of *q*_*sub*_ must follow a Hamiltonian cycle in *G*(*V*, *E*, *σ*). Accordingly, there is a Hamiltonian cycle in *G*′(*V*, *E*).

#### Corollary 2

*The problem “Can we perform ≤ d edits in graph G so that q will match in G?” is 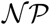-complete for |Σ| ≥* 2.

*Proof.* The setup used to prove Theorem 2 can be trivially extended to prove the above claim.

It is straightforward to prove that other problem variants, e.g., with linear gap penalty or affine gap penalty scoring functions are at least as hard as the edit-distance based formulations. Therefore, the sequence to graph alignment problem remains 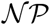-complete even on constant sized alphabets for these classes of scoring functions also if changes are permitted in the graph.

## 4 Sequence-to-Graph Alignment with Edits in Sequence

The sequence to graph alignment problem is polynomially solvable when changes are allowed on the query sequence alone [1, 20]. Here, we improve upon the state-of-the-art by presenting an algorithm with *O*(|*V*| + *m*|*E*|) run-time. Our algorithm matches the run-time complexity achieved previously by Rautiainen and Marschall [26] for edit distance, while improving that for linear and affine gap penalty functions. In addition, it is simpler to implement because it only uses elementary queue data structures. Note that edit distance is a special case of linear gap penalty when cost per unit length of the gap is 1, and substitution penalty is also 1. We first present our algorithm for the case of a linear gap penalty function, and subsequently show its generalization to affine gap penalty. From hereon, we assume that the sequence graph *G*(*V*, *E*, *σ*) is a character labeled graph, i.e., *σ*(*v*) ∈ *Σ*, *v* ∈ *V*. This assumption simplifies the description of the algorithm. Note that it is straightforward to transform a graph from string-labeled form to character-labeled form, and vice versa.

### 4.1 Linear Gap Penalty

#### Alignment Graph

In the literature on the classic sequence to sequence alignment problem, the problem is either formulated as a dynamic programming problem or an equivalent graph shortest-path problem in an appropriately constructed edge-weighted *edit graph* or *alignment graph* [18]. However, formulating the sequence to graph alignment problem as a dynamic programming recursion, while easy for directed acyclic graphs through the use of topological ordering, is difficult for general graphs due to the possibility of cycles. As it turns out, formulation as a shortest-path problem in an alignment graph is still rather convenient, even for graphs with cycles [1, 26]. The alignment graph, described below, is constructed using the given query sequence, the sequence graph and the scoring parameters.

The alignment graph is a weighted directed graph which is constructed such that each valid alignment of the query sequence to the sequence graph corresponds to a path from source vertex *s* to sink vertex *t* in the alignment graph, and vice versa (Figure 3). The alignment cost is equal to the corresponding path distance from the source to the sink. Note that the alignment graph is a multi-layer graph containing *m* ‘copies’ of the sequence graph, one in each layer. A column of dummy vertices is required in addition to accommodate the possibility of deleting a prefix of the query sequence. Edges that emanate from a vertex are equivalent to the choices available while solving the alignment problem. A formal definition of the alignment graph follows:

##### Definition 2.

*Alignment graph: Given a query sequence q, a sequence graph G*(*V, E, σ*)*, linear gap penalty parameters Δ_del_, Δ_ins_, and a substitution cost parameter Δ_sub_, the corresponding alignment graph is a weighted directed graph G_a_*(*V_a_, E_a_, ω_a_*)*, where V_a_* = ({1, …, *m*} × (*V ∪ {δ}*) *∪ {s, t} is the vertex set, and ω_a_*: *E_a_* → ℝ_≥0_ *is the weight function defined as*

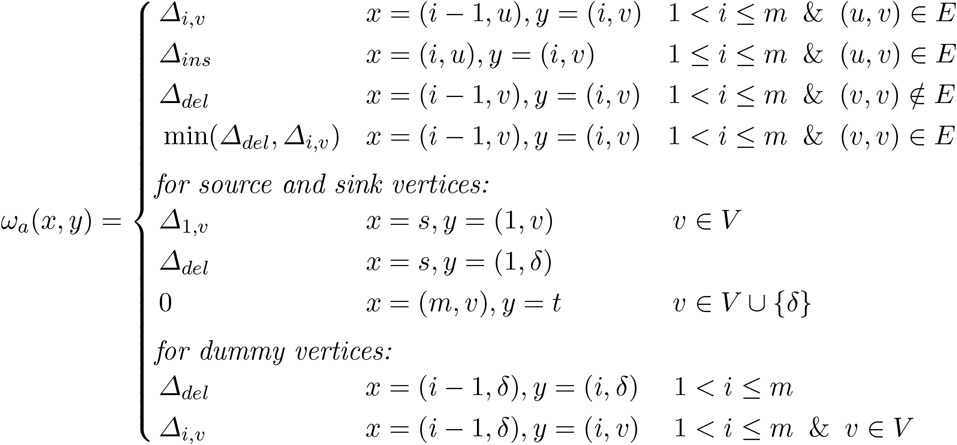

*Edges* (*x*, *y*) *∈ E_a_ are defined implicitly, as those pairs* (*x*, *y*) *for which ω_a_ is defined above. Δ_i,v_* = *Δ_sub_ if q*[*i*] ≠ *σ*(*v*)*, v ∈ V, and* 0 *otherwise. Δ_sub_ denotes the cost of substituting q*[*i*] *with σ*(*v*).

Existing definitions of the alignment graph [1, 26] did not include the dummy vertices, and were incomplete. Using the alignment graph, we reformulate the problem of computing an optimal alignment to finding the shortest path in the alignment graph. Even though the alignment graph defined by Amir *et al.* [1] has minor differences, proof in their work can be easily adapted to state the following claim:

**Fig. 3.**
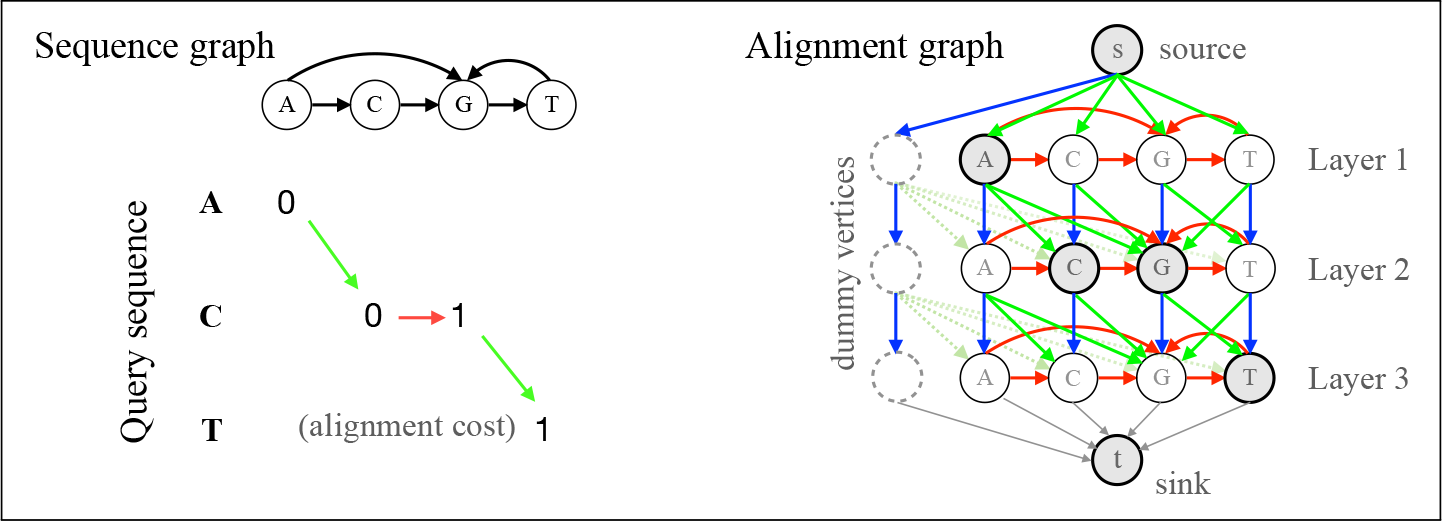
An example to illustrate the construction of an alignment graph from a given sequence graph and a query sequence. Multiple colors are used to show weighted edges of different categories in the alignment graph. The red, blue and green edges are weighted as insertion, deletion and substitution costs respectively.

##### Lemma 1 (Amir *et al.* [1]).

*Shortest distance from the source vertex s to the sink vertex t in the alignment graph G_a_*(*V_a_, E_a_, ω_a_*) *equals cost of optimal alignment between the query q and the sequence graph G*(*V, E, σ*).

One way of solving the above shortest path problem is to directly apply Dijkstra’s algorithm [1, 2]. However, it results in an *O* (*m*|*V*| log(*m*|*V*|)+ *m*|*E*| time algorithm. We next show how to solve the problem in *O*(|*V*| + *m*|*E*|) time.

##### Algorithm 1: Algorithm for sequence to graph alignment

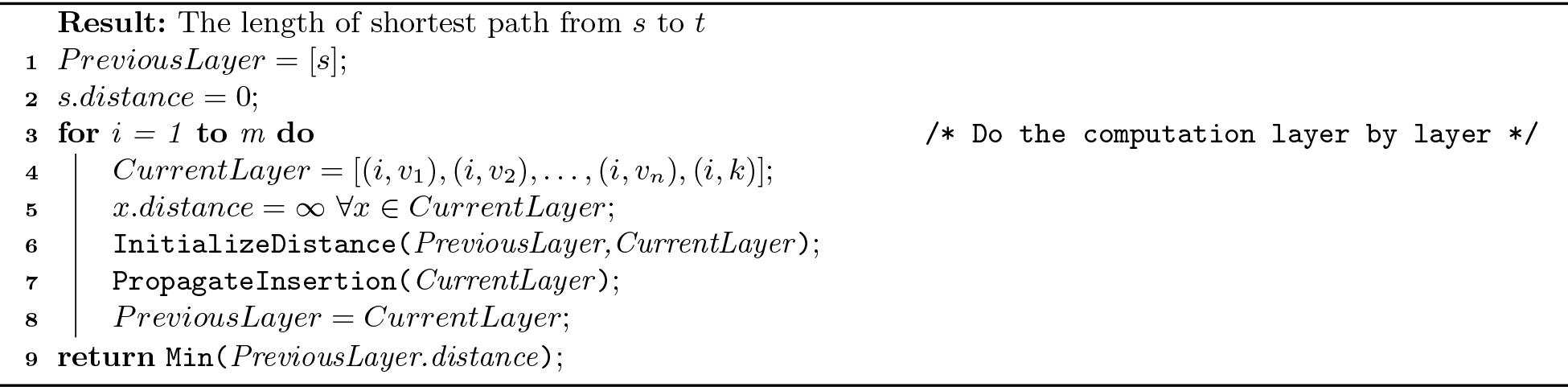

##### Algorithm 2: Algorithm to initialize and sort layer before insertion propagation

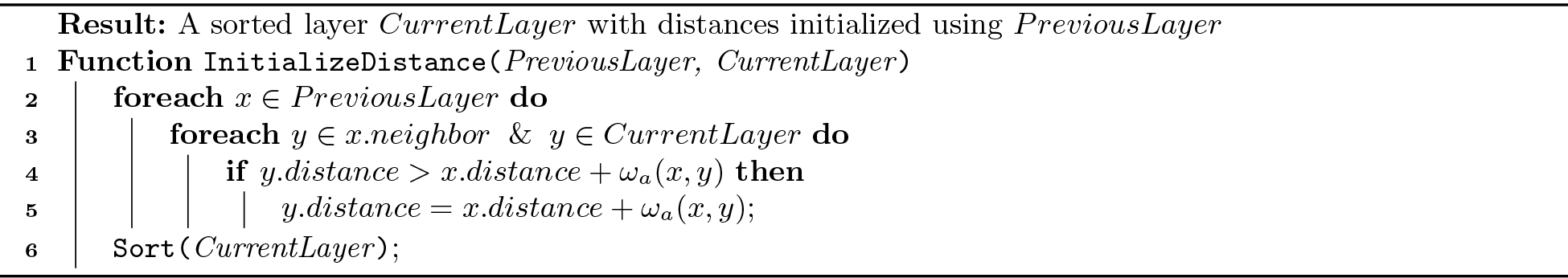

#### Proposed Algorithm

While searching for a shortest path from the source to the sink vertex, we compute the shortest distances from the source to intermediate vertices *V*_*a*_\{*s*, *t*} in the alignment graph. An edge from a vertex in layer *i* is either directed to a vertex in the same layer or to a vertex in the next layer. As a result, the shortest distances to nodes in a layer can be computed once the distances for the previous layer are known. This also makes it feasible to solve for the layers 1 to *m*, one by one [20]. We use a two-stage strategy to achieve linear *O*(|*V*| + |*E*|) run-time per layer. Before describing the details, we give an outline of the algorithm and its two stages.

Any path from the source vertex to a vertex *v* in a layer must extend a path ending in the previous layer using either a deletion or a substitution cost weighted edge. Afterwards, a path that ends in the same layer but not at *v* can be further extended to *v* using the insertion cost weighted edges if it results in the shortest path to the source. Roughly speaking, the first stage executes the former task, while the second takes care of the latter. The two stages together are invoked *m* times during the algorithm until the optimal distances are known for the last layer (Algorithm 1). Input to the first stage *InitializeDistance* is an array of the shortest distances of the vertices in previous layer sorted in non-decreasing order. This stage computes the ‘tentative’ distances of all vertices in the current layer because it ignores the insertion cost weighted edges during the computation. It outputs the sorted tentative distances as an input to the second stage *PropagateInsertion*. The PropagateInsertion stage returns the optimal distances of all vertices in the current layer while maintaining the sorted order for a subsequent iteration.

The following are two important aspects of our algorithm. First, we are able to maintain the sorted order of vertices by spending *O*(|*V*|) time per layer during the first stage (Lemma 2). Secondly, we propagate insertion costs through the edges in *O*(|*V*| + |*E*|) time per layer during the second stage by eluding the need for standard priority queue implementations (Lemmas 3-5). Both of these features exploit characteristics specific to the alignment graphs.

##### Algorithm 3: Algorithm to propagate insertions in the same layer

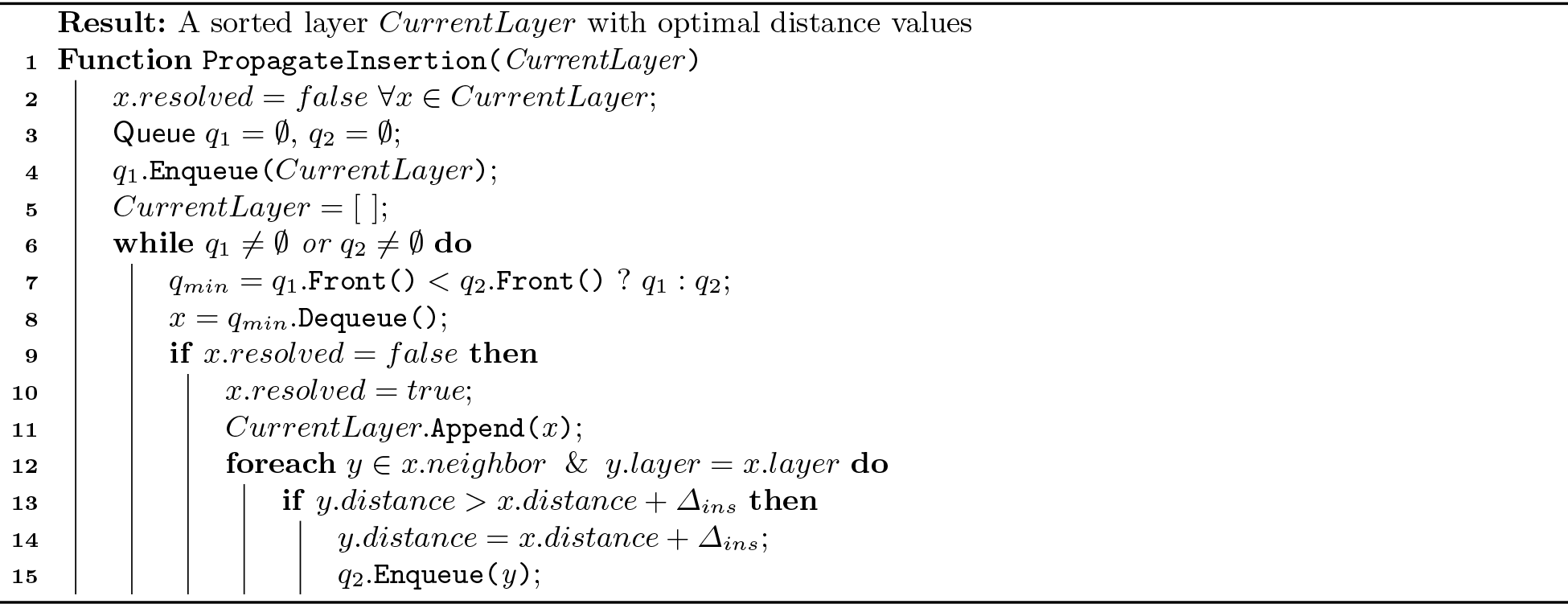

##### The InitializeDistance stage

We compute tentative distances for each vertex in the current layer by using shortest distances computed for the previous layer (Algorithm 2). Because all deletion and substitution cost weighted edges are directed from the previous layer towards the current, this only requires a straightforward linear *O*(|*V*| + |*E*|) time traversal (lines 2-5). In addition, we are required to maintain the current layer as per sorted order of distances. Note that vertices in the previous layer are already available in sorted order of their shortest distances from *s*. A vertex *v* in the previous layer can assign only three possible distance values (*v.distance*, *v.distance* + *Δ*_*sub*_, or *v.distance* + *Δ*_*del*_) to a neighbor in the current layer. By maintaining three separate lists for each of the three possibilities, we can create the three lists in sorted order and merge them in *O*(|*V*|) time. The relative order of vertices in the current layer can be easily determined in linear time by tracking the positions of their distance values in the merged list. As a result, the current layer can be obtained in sorted form in *O*(|*V*|) time and *O*(|*V*|) space, leading to the following claim.

###### Lemma 2.

*Time and space complexity of the sorting procedure in Algorithm 2 is O*(|*V*|).

##### The PropagateInsertion Stage

Note that the tentative distance computed for a vertex is sub-optimal if its shortest path from the source vertex traverses any insertion cost weighted edge in the current layer. One approach to compute optimal distance values is to process vertices in their sorted distance order (minimum first) and update the neighbor vertices, similar to Dijkstra’s algorithm. When processing vertex *v*, the distance of its neighbor should be adjusted such that it is no more than *v.distance* + *Δ*_*ins*_. Selecting vertices with minimum scores can be achieved using a standard priority queue implementation (e.g., Fibonacci heap); however, it would require *O*(|*E*| + |*V*| log |*V*|) time per layer. A key property that can be leveraged here is that all edges being considered in this stage have uniform weights (*Δ*_*ins*_). Therefore, we propose a simpler and faster algorithm using two First-In-First-Out queues (Algorithm 3). The first queue *q*_1_ is initialized with sorted vertices in the current layer, and the second queue *q*_2_ is initialized as empty (line 4). The minimum distance vertex is always dequeued from either of the two queues (line 8). As and when distance of a vertex is updated by its neighbor, it is enqueued to *q*_2_ (line 15). Following lemmas establish the correctness and an *O*(|*E*| + |*V*|) time bound for the PropagateInsertion stage in the algorithm.

###### Lemma 3.

*In each iteration at line 8, Algorithm 3 dequeues a vertex with the minimum overall distance in q*_1_ *and q*_2_.

*Proof.* The queue *q*_1_ always maintains its non-decreasing sorted order at the beginning of each loop iteration (line 6) in Algorithm 3 as we never enqueue new elements into *q*_1_. We prove by contradiction that *q*_2_ also maintains the order. Maintaining this invariant would immediately imply the above claim. Let *i* be the first iteration where *q*_2_ lost the order. Clearly *i* > 1. Because *i* is the first such iteration, new vertices (say *y*_1_, *y*_2_, …, *y*_*k*_) must have been enqueued to *q*_2_ in the previous iteration (line 15), and the vertex (say *x*) which caused these additions must have been dequeued (line 8). Note that the distance of all the new vertices, the *y*_*i*_’s, equals *x.distance* + *Δ*_*ins*_. Therefore, the vertex prior to *y*_1_ (say *y*_*pre*_) must have a distance higher than *y*_1_. However, this leads to a contradiction because if we consider the iteration when *y*_*pre*_ was enqueued to *q*_2_, the distance of the vertex that caused addition of *y*_*pre*_ could not be higher than the distance of the vertex *x*.

###### Lemma 4.

*Once a vertex is dequeued in Algorithm 3, its computed distance equals the shortest distance from the source vertex*.

*Proof.* Lemma 3 establishes that Algorithm 3 processes all vertices that belong to the current layer in sorted order. Therefore, it mimics the choices made by Dijkstra’s algorithm [5].

###### Lemma 5.

*Algorithm 3 uses O*(|*V*| + |*E*|) *time and O*(|*V*|) *space to compute shortest distances in a layer*.

*Proof.* Each vertex in the current layer enqueues its updated neighbor vertices into *q*_2_ at most once. Note that distance of a vertex can be updated at most once, therefore the maximum number of enqueue operations into *q*_2_ is |*V*|. In addition, enqueue operations are never performed in *q*_1_. Accordingly, the number of outer loop iterations (line 6) is bounded by *O*(|*V*|). The inner loop (line 12) is executed at most once per vertex, therefore the amortized run-time of the inner loop is *O*(|*V*| + |*E*|).

The above claims yield an *O*(*m*(|*V*| + |*E*|)) time algorithm for aligning the query sequence to sequence graph. Assuming a constant alphabet, we can further tighten the bound to *O*(|*V*| + *m*|*E*|) by using a simple preprocessing step suggested in [26]. This step transforms the sequence graph by merging all vertices with 0 in-degree into ≤ |*Σ*| vertices. As a result, the preprocessing ensures that the count of vertices in the new graph is no more than |*E*| + |*Σ*| without affecting the correctness. Summary of the above claims is presented as a following theorem:

###### Theorem 3

*Algorithm 1 computes the optimal cost of aligning a query sequence of length m to graph G*(*V, E, σ*) *in O*(|*V*| + *m*|*E*|) *time and O*(|*V*|) *space using a linear gap penalty cost function*.

It is natural to wonder whether there exist faster algorithms for solving the sequence to graph alignment problem. As noted in [26], the sequence to sequence alignment problem is a special case of the sequence to graph alignment problem because a sequence can be represented as a directed chain graph with character labels. As a result, existence of either *O*(*m*^1−*∊*^|*E*|) or *O*(*m*|*E*|^1−*∊*^), *∊* > 0 time algorithm for solving the sequence to graph alignment problem is unlikely because it would also yield a strongly sub-quadratic algorithm for solving the sequence to sequence alignment problem, further contradicting SETH [3].

### 4.2 Affine Gap Penalty

Supporting affine gap penalty functions in the dynamic programming algorithm for sequence to sequence alignment is typically done by using three rather than one scoring matrix [10]. Similarly, the alignment graph can be extended to contain three sub-graphs with substitution, deletion, and insertion cost weighted edges respectively [26]. The edge weights are adjusted for the affine gap penalty model such that a cost for opening a gap is penalized whenever a path leaves the match sub-graph to either the insertion or the deletion sub-graph (Appendix Figure 4). The properties that were leveraged to design faster algorithm for linear gap penalty functions continue to hold in the new alignment graph. In particular, the sorting still requires linear time during the InitializeDistance stage, and insertion propagation is still executed over uniformly weighted edges in the insertion sub-graph. As a result, the two-stage algorithm can be extended to operate using affine gap penalty function in the same time and space complexity as with the linear gap penalty function.

**Fig. 4.**
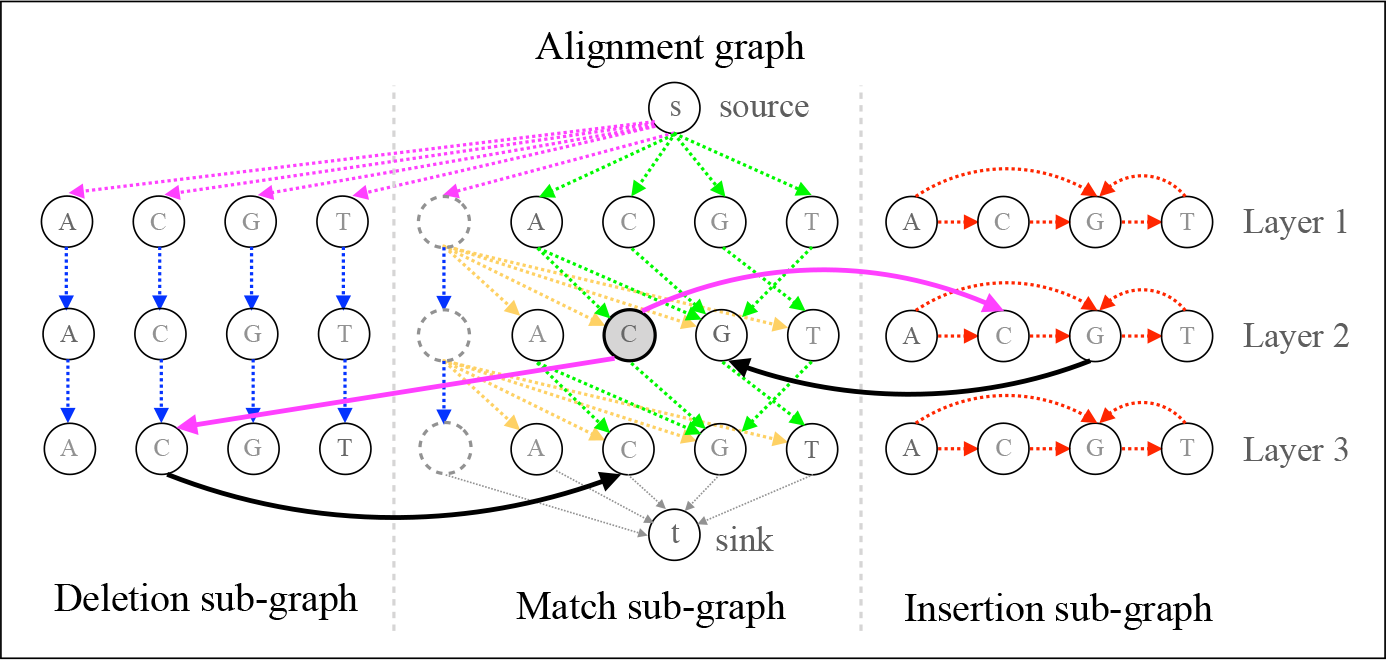
An example to illustrate the construction of an alignment graph for sequence to graph alignment using affine gap penalty. The alignment graph now contains three sub-graphs separated by the gray dash lines. The deletion and insertion weighted edges in the alignment graph for linear gap penalty are shifted to deletion sub-graph and insertion sub-graph, respectively. Their weights are also changed to the gap extension penalty. Besides, more edges are added to connect the sub-graphs with each other. For simplicity, we use the highlighted vertex as an example to illustrate how to open a gap and extend it. The weight of magenta colored edges is the sum of gap open penalty and gap extension penalty, and the weight of the black colored edges is 0.

## 5 Conclusions and Open Problems

In this paper, we show that the sequence to graph alignment problem is 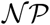-complete when changes are allowed in the sequence graph, for any alphabet of size ≥ 2. When changes are allowed in the query sequence alone, we provide a faster polynomial time algorithm that generalizes to linear gap penalty and affine gap penalty functions. The proposed algorithms use elementary data structures, therefore are simple to implement. Overall, the theoretical results presented in this work enhance the fundamental understanding of the problem, and will aid the development of faster tools for mapping to graphs. The alignment problem for sequence graphs is a rich area with several unsolved problems. For the intractable problem variants, development of faster exact and approximate algorithms are fertile grounds for future research. In addition, working towards robust indexing schemes and heuristics that scale to large input graphs and different sequencing technologies remains an important direction.

## Acknowledgements

This work is supported in part by US National Science Foundation grant CCF-1816027. Yu Gao was supported by the ACO Program at Georgia Institute of Technology.

## 6 Appendix

